# BIDS Apps: Improving ease of use, accessibility, and reproducibility of neuroimaging data analysis methods

**DOI:** 10.1101/079145

**Authors:** Krzysztof J. Gorgolewski, Fidel Alfaro-Almagro, Tibor Auer, Pierre Bellec, Mihai Capotă, M. Mallar Chakravarty, Nathan W. Churchill, Alexander Li Cohen, R. Cameron Craddock, Gabriel A. Devenyi, Anders Eklund, Oscar Esteban, Guillaume Flandin, Satrajit S. Ghosh, J. Swaroop Guntupalli, Mark Jenkinson, Anisha Keshavan, Gregory Kiar, Franziskus Liem, Pradeep Reddy Raamana, David Raffelt, Christopher J. Steele, Pierre-Olivier Quirion, Robert E. Smith, Stephen C. Strother, Gaël Varoquaux, Tal Yarkoni, Yida Wang, Russell A. Poldrack

## Abstract

The rate of progress in human neurosciences is limited by the inability to easily apply a wide range of analysis methods to the plethora of different datasets acquired in labs around the world. In this work, we introduce a framework for creating, testing, versioning and archiving portable applications for analyzing neuroimaging data organized and described in compliance with the Brain Imaging Data Structure (BIDS). The portability of these applications (BIDS Apps) is achieved by using container technologies that encapsulate all binary and other dependencies in one convenient package. BIDS Apps run on all three major operating systems with no need for complex setup and configuration and thanks to the comprehensiveness richness of the BIDS standard they require little manual user input. Previous containerized data processing solutions were limited to single user environments and not compatible with most multi-tenant High Performance Computing systems. BIDS Apps overcome this limitation by taking advantage of the Singularity container technology. As a proof of concept, this work is accompanied by 22 ready to use BIDS Apps, packaging a diverse set of commonly used neuroimaging algorithms.

**Author Summary:** Magnetic Resonance Imaging (MRI) is a non-invasive way to measure human brain structure and activity that has been used for over 25 years. There are thousands MRI studies performed every year generating a substantial amount of data. At the same time, many new data analysis methods are being developed every year. The potential of using new analysis methods on the variety of existing and newly acquired data is hindered by difficulties in software deployment and lack of support for standardized input data. Here we propose to use container technology to make deployment of a wide range of data analysis techniques easy. In addition, we adapt the existing data analysis tools to interface with data organized in a standardized way. We hope that this approach will enable researchers to access a wider range of methods when analyzing their data which will lead to accelerated progress in human neuroscience.

## Introduction

The last 25 years have witnessed a proliferation of methods for imaging the human brain (including structural, diffusion and functional Magnetic Resonance Imaging, Positron Emission Tomography, Electroencephalography and Magnetoencephalography). These methods have been accompanied by literally thousands of different algorithms for signal denoising, normalization, feature extraction, and statistical analysis. Modern analysis pipelines for neuroimaging often consist of dozens of steps and rely on software developed by multiple external groups with each group often developing their own idiosyncratic parameter settings even when using the same software packages. The increasing complexity of neuroimaging data analysis has led to many discoveries, and the flexibility provided by the plethora of feature extraction methods has allowed cognitive and clinical neuroscientists to develop new theories about the relationships between brain and behavior in healthy and diseased populations.

However, due to the intrinsic heterogeneity of scientific software (arising from the fact that it rarely is developed for widespread distribution), installing, configuring, and running many of the available methods is often difficult. Most neuroimaging software packages run natively on Linux [1] and (to a lesser extent) Mac OS X; however, Windows users often cannot run software natively. Operating system aside, many scientific packages depend on external libraries (often requiring a particular, sometimes outdated version), and require complex configurations of environment variables and/or data files.

There have been some attempts in the field of neuroinformatics to solve this issue. The most notable is the NeuroDebian project [2], which provides Debian and Ubuntu Linux distributions with packages containing many of the popular neuroimaging software tools. Installing packages prepared by the NeuroDebian team is very easy, as it can be performed with the built-in Debian/Ubuntu package management system. However, this solution only applies to Linux systems, as Mac OS X and Windows users are required to install Linux inside a Virtual Machine (VM). Additionally, creating new Debian/Ubuntu packages is a non-trivial task, which may limit the rate at which new software is added to NeuroDebian.

Another consequence of these deployment and installation issues is that they make it very difficult to perfectly recreate analysis pipelines, which exacerbates reproducibility issues in neuroimaging [3]. This is partly due to journal space limitations, which typically preclude a full accounting of the scientific software stack used to generate results. But even knowing the versions of all of the software tools employed in an analysis is rarely sufficient to completely reproduce a workflow. Studies have shown that operating system type, version, and even hardware architecture can influence results in a significant manner [4–6]. Such issues are troubling in and of themselves and call for thorough investigations into the sources of this variability. They also raise serious practical concerns for extended longitudinal studies that need to maintain the same software (and sometimes even hardware) stack along the time span of an experiment (e.g. a software upgrade midway through the analysis could lead to spurious differences between groups processed at different points in time).

One potential solution to ease the deployment problem proposed in the bioinformatics community is to create Virtual Machines (VMs) capturing all of the necessary dependencies for a workflow [7, 8]. Running a Virtual Machine does, however, come with significant performance overhead; furthermore, only a few High Performance Computing (HPC) systems (which have become the primary computational resource for many academics in the last decade) allow their users to run VMs on large clusters. Building on the virtual machine concept, a more lightweight solution has recently become more prevalent in the industry, known as ‘operating-system-level-virtualization’ or (more commonly) *containers*. In contrast to VMs, containers share the same kernel with the host operating system and thus deliver much better performance. The bioinformatics community has once again been at the forefront of the adoption of this new technology, with the aim of improving research reproducibility [9–12]. Most proposed solutions have been based on a particular implementation of the container concept: ‘Docker’, which, due to its kernel and security requirements, is difficult or impossible to use in a multi-tenant (serving multiple unprivileged users) environment such as an HPC system (which is often the most cost-effective computational resource available to researchers).

Finally, most existing neuroimaging data processing pipelines expect input datasets to be organized and described in different and idiosyncratic ways. To account for variability in input data organization and a lack of consensus in terms of metadata description, data processing workflows often require users to input metadata manually - in a different fashion for each pipeline. This expandable step can cause errors and lead to incorrect results, while also making it harder to integrate processing pipelines into automated analysis platforms such as those developed to provide “Science as a Service” [13].

After careful evaluation of existing solutions for the specific problems faced by the neuroimaging community, we have developed a framework for sharing and executing neuroimaging analysis pipelines that improve ease of use, accessibility, and reproducibility for users of all three major operating systems, as well as for researchers using HPC or cloud computing systems. The framework overcomes Docker’s inability to run on multi-tenant HPC systems by capitalizing on the Singularity [14] container technology developed specifically for HPC use. To minimize the number of manual inputs required from researchers--and hence reduce the number of errors arising from misinterpretation of those inputs--the framework capitalizes on the recently introduced Brain Imaging Data Structure (BIDS) standard [15] for organizing and describing datasets. Correspondingly, the proposed framework is named BIDS Apps. We describe here the anatomy of a BIDS App, the infrastructure used for building, testing and archiving BIDS Apps, as well as steps necessary to run BIDS Apps in various scenarios.

## Results

### What is a BIDS App?

A BIDS App is a container image capturing a neuroimaging pipeline that takes a BIDS-formatted dataset as input. Since the input is a whole dataset, apps are able to combine multiple modalities, sessions, and/or subjects, but at the same time need to implement ways to query input datasets. Each BIDS App has the same core set of command-line arguments, making them easy to run and integrate into automated platforms. BIDS Apps are constructed in a way that does not depend on any software outside of the container image other than the container engine.

BIDS Apps rely upon two technologies for container computing:

1. Docker - for building, hosting as well as running containers on local hardware (running Windows, Mac OS X or Linux) or in the cloud.
2. Singularity - for running containers on HPCs [14].

BIDS Apps are deposited in the Docker Hub (http://hub.docker.com) repository, making them openly accessible. Each app is versioned and all of the historical versions are available to download. By reporting the BIDS App name and version in a manuscript, authors can provide others with the ability to exactly replicate their analysis workflow.

Docker is used for its excellent documentation, maturity, and the Docker Hub service for storage and distribution of the images. Docker containers are easily run on personal computers and cloud services. However, the Docker Engine was originally designed to run different components of web services (HTTP servers, databases etc.) using cloud resources. Docker thus requires root or root-like permissions, as well as modern versions of Linux kernel (to perform user mapping and management of network resources); though this is not a problem in context of renting cloud resources (which are not shared with other users), it makes it difficult or impossible to use in a multi-tenant environment such as an HPC system, which is often the most cost-effective computational resource available to researchers. Singularity, on the other hand, is a unique container technology designed from the ground up with the encapsulation of binary dependencies and HPC use in mind. Its main advantage over Docker is that it does not require root access for container execution and thus is safe to use on multi-tenant systems. In addition, it does not require recent Linux kernel functionalities (such as namespaces, cgroups and capabilities), making it easy to install on legacy systems.

**Figure 1.**
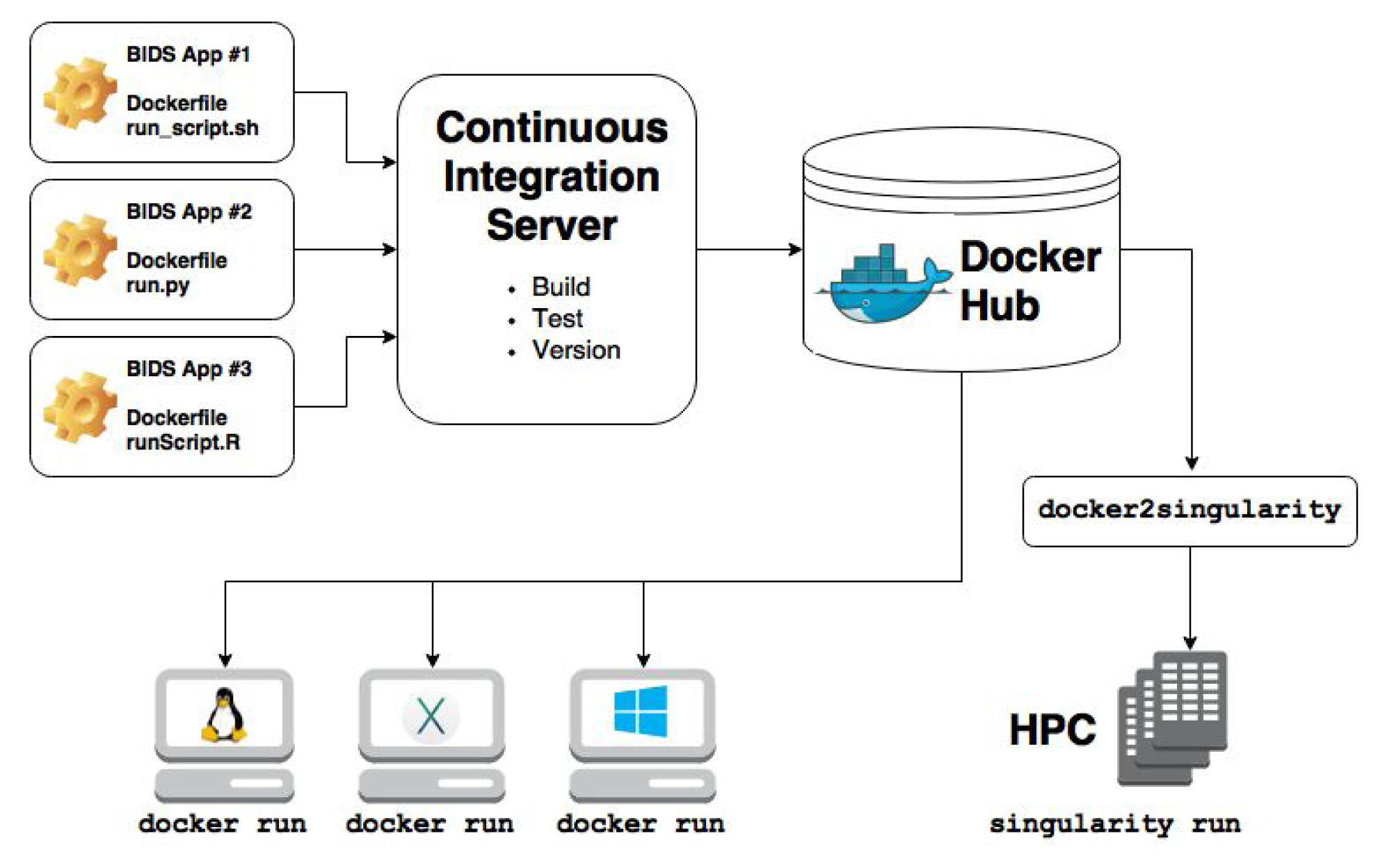
Overview of the creation and use of BIDS Apps. The source code of each App is stored in separate GitHub repository. Each repository is connected to a Continuous Integration server responsible for building testing and deploying the corresponding App. For every new release of an App, a new container image is deposited in Docker Hub. Users can directly download and run the BIDS Apps container images either directly using Docker on any Windows, Mac, and Linux machine or convert them to Singularity and run them on an HPC.

### How to create a BIDS App?

#### BIDS Apps Forge

Inspired by the conda-forge project (https://conda-forge.github.io/), BIDS Apps development is centered around a GitHub organization (http://github.com/BIDS-Apps) that maintains a code repository for each BIDS App (see Figure 1). Every repository hosts a Dockerfile describing how to build the container image, a lightweight wrapper for providing a unified command-line interface as well as parsing the BIDS input, and brief documentation. BIDS Apps are an evolution of lightweight wrappers such as the Interface class developed within Nipype [16], with the added advantage of being programming language agnostic. Since BIDS Apps merely serve the purpose of capturing dependencies and providing a unified way of calling the relevant program (through the command-line interface), the repositories in the github.com/BIDS-Apps organization do not actually host the source code or data of the pipelines and workflows being wrapped: these are contained in the built Docker image stored on DockerHub. Building a BIDS App starts with creating a new repository and populating it with a Dockerfile and a run script.

#### Dockerfile creation

A *Dockerfile* is a script written in a domain specific language that describes the steps necessary to build a Docker image. The Docker project provides excellent documentation and a tutorial on how to write Dockerfiles (https://docs.docker.com/engine/reference/builder/). However, because the container images built in the BIDS-Apps format are ultimately intended to alternatively run under Singularity, there are some additional requirements that must be followed:

**•** Apps cannot rely on having elevated permissions inside the container image (in contrast to Docker, processes inside a Singularity container run with the privileges of the user running the container).

**•** Environment variables must be set using the ENV statement within the relevant Dockerfile, rather than relying on config files (such as /root/.bashrc).

**•** Apps should not write anywhere outside of /tmp, $HOME, and the specific output folder provided as a command-line argument.

To facilitate the process of writing Dockerfiles, we have created a set of templates that include installation steps for the most popular neuroimaging tools (FSL, FreeSurfer, AFNI, ANTs etc.): https://github.com/BIDS-Apps/dockerfile-templates. Those templates are also available as container images that can be used directly as a base for new BIDS Apps Dockerfiles using the FROM statement.

#### Command-line interface

To improve user experience and ability to integrate BIDS Apps into various computational platforms, each App follows a set of core command-line arguments:

~~~
runscript input_dataset output_folder analysis_level
~~~

For example:

~~~
runscript /data/ds114 /scratch/outputs participant
~~~

- `input_dataset` provides a path to the dataset to be analyzed (read-only), which must conform to the BIDS standard
- `output_folder` is the folder where results of the analysis will be stored
- `analysis_level` denotes the stage of the analysis that will be performed

To facilitate easy and efficient execution, analyses in BIDS Apps can be split into stages (see Materials and Methods). In the simplest design, an App would run in two stages: ‘participant’ and ‘group’. The ‘participant’ stage runs a first level analysis that can be performed independently for each subject in the dataset (and thus can be executed in parallel). The analysis may optionally be restricted to a subset of participants using the --`participant_label` argument. The group level analysis runs on the outputs from the participant level analysis and cannot be split into independent parallel jobs. This scheme is inspired by the MapReduce programming model [17]. Multiple such MapReduce steps can be defined for a single pipeline (full specification of the command-line scheme can be found in the Supplementary Materials).

There are no restrictions on what language is used to write the wrapper script as long as it conforms to the prescribed interface. However, to make it easier to generate new BIDS Apps we have created a basic implementation in Python that can be imported into a new script and filled with App-specific options: https://github.com/BIDS-Apps/bidscmd. We also provide several utilities that make it easier to work with BIDS-compatible directory structures--most notably, the PyBIDS Python package (https://github.com/INCF/pybids), which provides tools for simple but powerful logical queries over entities defined in the BIDS specification (e.g., retrieving a list of all unique subjects; getting the fieldmap files for all subjects with a valid first scanning run; etc.).

In addition to conforming to a standardized command-line argument scheme, run scripts are also responsible for validation of the input data before running any analysis. To facilitate the process we have developed a command-line validator that checks whether the input datasets are compliant with the BIDS standard https://github.com/INCF/bids-validator. Because not all BIDS compatible datasets can be analyzed by all BIDS Apps (e.g. a surface reconstruction pipeline requires a high-resolution T1 weighted image), the validator can be configured to reject datasets with particular properties. Integrating the validator is as easy as calling an external command; Dockerfiles and container images with the validator pre-installed are also available. BIDS Apps developers can also choose to implement validation steps themselves if the requirements of their pipelines cannot be easily checked by the standard validator.

#### Building and testing container images

For each BIDS App a Docker image is built and run on a set of lightweight example BIDS datasets [18]. This execution tests that the command-line interface and BIDS support are correctly implemented (each BIDS App is required to include at least one smoke or integration test running on an example BIDS dataset). The tests are run forcing read-only containers, to ensure compatibility with Singularity (which imposes read-only mode). If the Docker image builds successfully and passes all tests, it is assigned a unique version (based on the tag obtained from the corresponding GitHub repository) and uploaded to Docker Hub with a version tag. Since version tags are unique and Docker images stored on Docker Hub are never overwritten, all historical versions of each BIDS App will always be accessible. Building, testing and archiving of BIDS Apps is performed automatically through a continuous integration service (CircleCI). Automation of testing and versioning improves reliability due to minimization of human errors.

To facilitate the process of creating new BIDS Apps we have made an example App that can also be used as a template for new Apps: https://github.com/BIDS-Apps/example. Additional documentation and tutorials are available at http://bids-apps.neuroimaging.io. Developers seeking help are also encouraged to subscribe to the bids-app-dev mailing list at https://groups.google.com/d/forum/bids-apps-dev.

## Available BIDS Apps

At the moment there are 22 BIDS Apps (see Table 1) in the repository, most of which were developed by participants in the 2016 sprint (see Materials and Methods). The Apps span different imaging modalities (structural, functional and diffusion MRI) as well as languages (Python, C++, MATLAB/Octave, OpenCL).

**Table 1.**
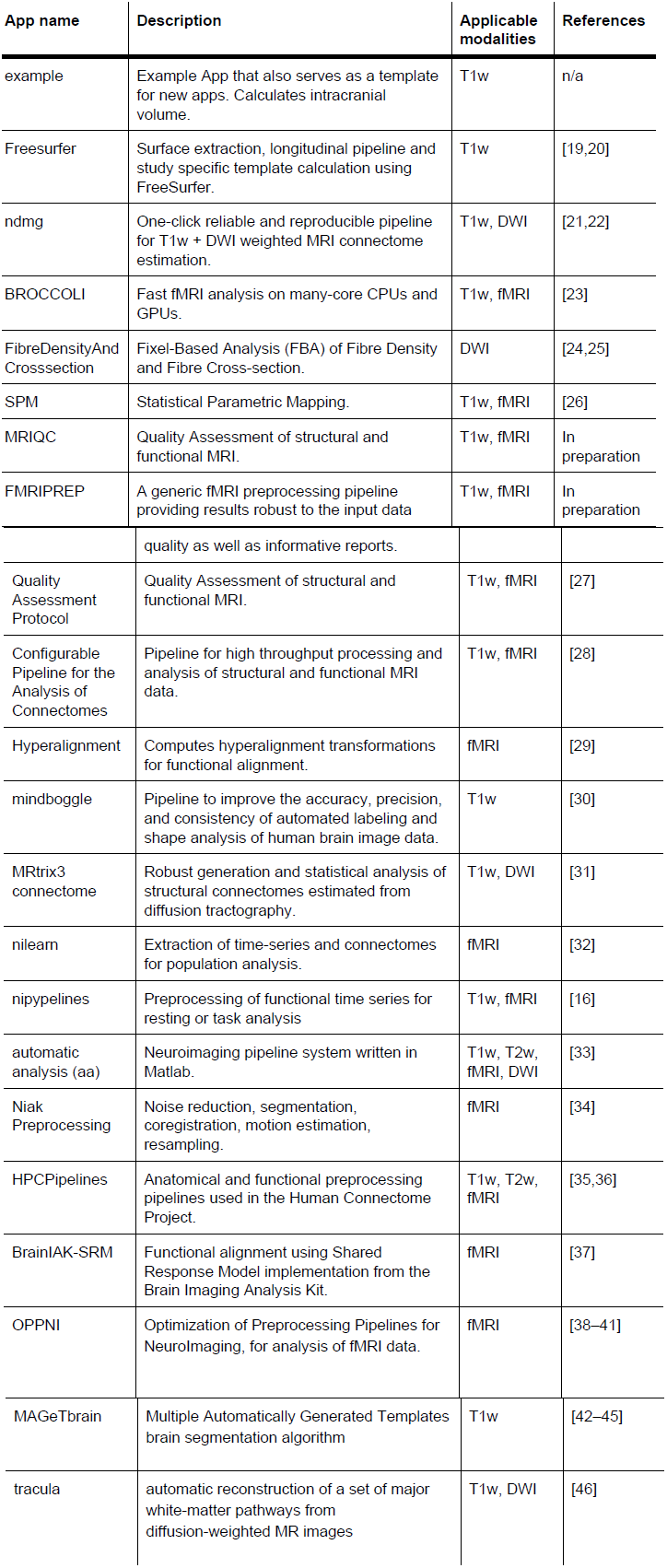
List of currently available BIDS Apps. 19–46.

## Running a BIDS App locally

Running a BIDS App on a local system can be performed using Docker, which is easy to install on all three major operating systems. To run the first stage of the example BIDS App for participant number 01 the user needs to open a console (terminal or cmd) and type:

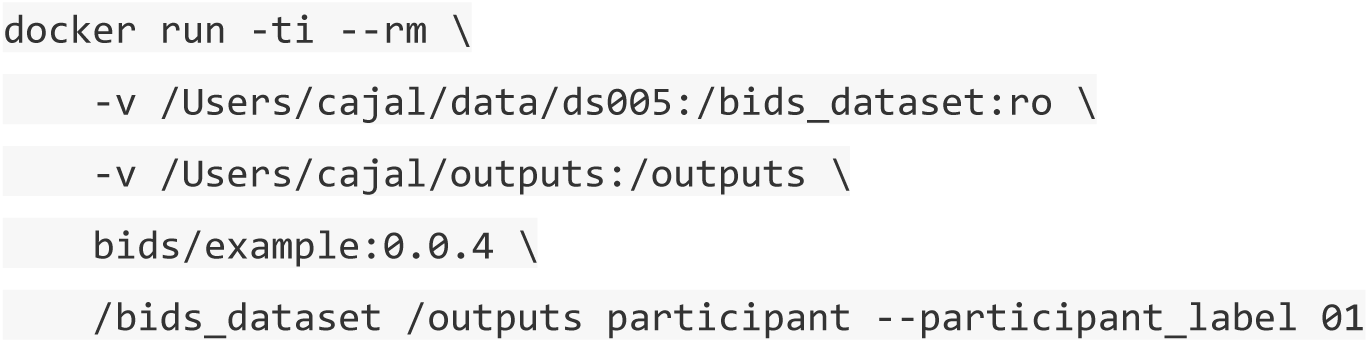

Where /Users/cajal/data/ds005 is the path to the input dataset and /Users/cajal/outputs the path where results should be stored. If the BIDS App was not run before on this machine, the Docker image will be automatically downloaded from the Docker Hub.

## Running a BIDS App on a cluster (HPC)

On many academic clusters, Singularity can be used to run containers (due to its minimal dependencies and security concerns Singularity is more likely to be approved for multi-tenant systems usage than Docker). In these setting, to run a BIDS App, it first needs to be saved to a Singularity-compatible image file. This step needs to be performed outside of the cluster (for example on a laptop) and requires Docker:

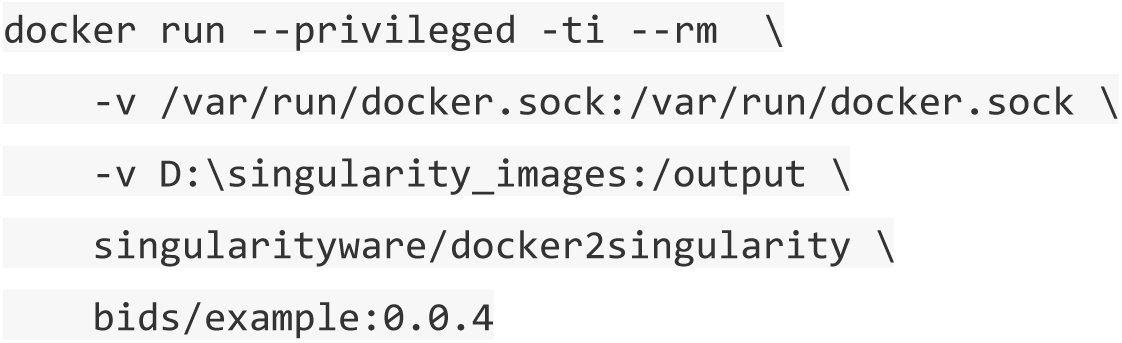

Where D:\singularity_images is a path where the Singularity image will be stored. After transferring the .img file to a cluster it can be run like any other executable:

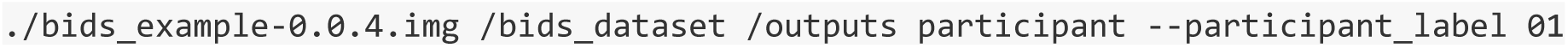

It is worth noting that copying and running Singularity images does not require elevated permissions.

## Discussion

We have proposed a new way of distributing easy-to-use, reproducible neuroimaging analysis workflows that can run on all three major operating systems as well as multi-tenant clusters. Each BIDS App encapsulates all of its binary dependencies providing the means for reproducible analysis as well as an ultimate source of provenance information. Due to the BIDS standard used for the organization of input data, errors caused by manually provided metadata are minimized. Finally, the unified command-line interface structure combined with flexible MapReduce-style execution schemes lends BIDS Apps to easy integration into data analysis platforms as well as efficient execution on computational clusters independently of the particular scheduling software.

To prove the viability of the BIDS Apps concept we have developed 20 Apps representing a diverse set of neuroimaging software originating from many different labs. We are expecting that the number of available apps will grow in the future. We are actively encouraging neuroimaging methods developers to deploy their tools as BIDS App to further grow the library of available pipelines.

One of the goals of this project was to deliver software that could be applied to wide range of datasets without the need for much user interaction. It is important to emphasize that this goal is not at odds with the ability to modify various parameters of the workflows encapsulated in BIDS Apps. This is an important ability since many researchers have unique needs and many data analysis decisions are considered open scientific questions. The ability to modify internal parameters vary across the presented BIDS Apps, but many of them provide it either in form of command line parameters or input config files. We hope that with the help of new users app developers will be able to learn which parameters should be exposed for modification.

It is also important to note that fact that BIDS Apps framework even though is built on container technology (which can be misinterpreted as a “black box”) it does not imply a particular software development philosophy. Some BIDS Apps are interfaces to open source projects with vibrant developer communities built around public developer interactions while some are developed in a restricted access setting by a smaller set of developers located at a particular institute. By choosing GitHub as the platform for hosting BIDS Apps we hope to foster a vibrant and open community of developers contributing to run scripts, Dockerfiles, and documentation.

Even though similar solutions based on Docker have been proposed in the past, none have addressed the problem of running Docker containers on HPCs. One study evaluating Docker in the context of HPC use [47] had to limit access to the Docker-enabled cluster only to “trusted” users due to security concerns (personal communication). The solution proposed here combines mature container building tools provided for multiple operating systems by the Docker project with HPC compatibility through Singularity.

Additionally, similarly to previous proposals [9,10], the presented solution puts a strong emphasis on clear versioning of container images and the ability to access all previous versions. We envisage that this feature will be very valuable in the context of longitudinal studies (where the same software stack needs to be used over many years) as well as for accurately reporting a set of computational methods in a publication and later replicating them (which can be achieved by simply referencing a BIDS App with a corresponding version). Strict versioning of BIDS Apps is achieved by careful management of Docker images on Docker Hub via a Continuous Integration service which also is responsible for testing the Docker images, further reducing potential errors.

It is also worth noting that even though containers increase the reproducibility of scientific results, they do not solve the problem of sensitivity of some results to different operating systems, architectures or third party libraries. In particular, numerical and statistical instabilities create an artificial dependence of results on hardware details that can only be addressed algorithmically. Further research is necessary to assess the robustness of published results to these factors. BIDS Apps can help in this endeavor to a certain extent - for example, a single analysis can be run using different versions of the same BIDS App to see the variance in results (different flavors of the same App using different Linux distributions could be created for this purpose). However this approach has limits, further work is needed to better understand the potential for variance in results due to different Linux kernel versions across different systems and hardware architectures since containers do not encapsulate this.

Another advantage of container-based solutions is that the user can run software in an almost identical software ecosystem as the one used for its development. This reduces the number of problems experienced by users due to nuanced differences in system configuration, such as system libraries or software versions. It also makes maintaining the software significantly easier for the developers, who do not need to support a variety of different configurations and can easily reproduce errors.

Even though BIDS Apps as a form of neuroimaging software distribution scheme can be perceived as performing a similar task as the NeuroDebian project, the two initiatives, in fact, complement each other. Many of the example BIDS Apps presented in this manuscript use Debian as their base distribution and benefit from the ease of installation provided by the NeuroDebian project. It not only makes Dockerfiles shorter and easier to maintain but due to the network of NeuroDebian mirror servers the build process is more reliable than downloading software from their original locations. On the other hand, the NeuroDebian project benefits from the BIDS App distribution scheme by exposing software previously limited only to Debian based distribution to all flavors of Linux. This is important considering that many HPCs run on RedHat and CentOS Linux distributions rather than Debian.

Future work will involve engaging more developers of neuroimaging methods to create BIDS Apps. Additionally, there are plans for developing a repository that would be independent of Docker Hub to provide Open Container Consortium as well as Singularity-compatible images. This would provide improved sustainability of the project, but also remove the conversion step (from Docker to Singularity) currently necessary for running BIDS Apps on clusters.

Work is currently underway to facilitate the integration of BIDS Apps in other platforms such as XNAT [48], CBRAIN [49], or cloud-based services like Amazon Web Service (AWS). The apps will include a machine-readable description of input arguments, their descriptions and acceptable values (based on the Boutiques application descriptor [50]. Another benefit of such a description is to allow developers integrating BIDS Apps into their platforms to automatically generate user interfaces, and to improve validation of input parameters.

Singularity is not the only solution proposed to handle containers on HPCs. “Shifter” (as of September 2016 only available as pre-release beta version) has been discussed in the context of running containerized academic software [51]. The principles behind Shifter are similar to Singularity, but Shifter also attempts to tackle the problem of managing the container images. Because of this, it depends on several services (Redis and MongoDB servers, worker processes, etc.) that make setting it up and maintaining it more involved than Singularity. However, despite the differences, all BIDS Apps would be able to run on clusters running Shifter.

Singularity is despite its many advantages is not perfect. The need to convert Docker container images outside using docker2singularity tool is less than ideal. In the future, this will be a new feature of Singularity (currently in development) that will allow HPC users to directly import container images from Docker Hub (without the requirement of elevated privileges or access to Docker software). As mentioned above Singularity due to its minimal system requirements is easy to install. We are currently aware of 51 HPCs spread around the world that support Singularity (https://docs.google.com/spreadsheets/d/1Vc_1prq_1WHGf0LWtpUBY-tfKdLLM_TErjnCe1mY5m0/pub?gid=1407658660). This list includes systems located at prestigious institutions such as National Institutes of Health, Massachusetts Institute of Technology, and Stanford University as well as the 17th most powerful HPC in the world - Stampede, located at Texas Advanced Computing Center (https://www.top500.org/system/177931). Nonetheless, Singularity is still fairly new and it is fair to assume that it is not installed on most existing HPCs. Finally, Singularity container images require root permission to create or modify (this is why conversion from Docker needs to be performed outside of the target HCP). On one side this can be perceived as inconvenience preventing users from performing quick modifications of container images. However, on of the other side, the side effect of this limitation is that users are forced to create a new container image file each time an app is updated which promotes versioning and reproducibility.

The proposed way of archiving BIDS Apps - using Docker Hub - even though convenient and cost effective (using Docker Hub is free for public images) might pose challenges in the long term. Docker Hub does not make any commitments about the availability of images. To avoid potential loss of images uploaded to Docker Hub we plan to periodically backup all of the BIDS Apps in a repository dedicated for long term academic storage such as the Stanford Digital Library.

Despite its relatively young age, we are already seeing interest in various BIDS Apps. Some of the early adopters helped resolve issues and fix bugs. According to Docker Hub, BIDS Apps have been so far been downloaded 1389 times. Even though it will take up to two years until we start seeing papers describing studies that were using BIDS Apps, we already know of two studies that used BIDS Apps for data preprocessing (personal communication).

In the future, we plan to extend the BIDS Apps framework by adding a specification for analysis outputs and extending the input specification beyond MRI data. An effort is currently underway to define BIDS Derivatives specification that would cover description and organization of data analysis outputs. Adoption of such standard will allow chaining multiple apps. The FMRIPREP app already supports an early version of the BIDS Derivatives specification. In a similar vein, two extensions of the BIDS standard are in final stages of development: BIDS MEG and BIDS PET. That effort - when finalized - will allow the BIDS Apps family to be extended by tools dedicated to modalities other than MRI.

It is also worth mentioning that even though the proposed framework is focused on the analysis of neuroimaging data, a similar scheme could be applied to other types of data. The only elements that are specific to human neuroimaging in BIDS Apps are the format of the input data and the naming convention for command line parameters (“participant” and “group”). Other fields with well-established data standards can easily adopt the same way of constructing, testing and archiving pipelines with their software dependencies.

BIDS Apps is a framework to help neuroimaging practitioners deploy the complex data processing workflows that they require in their research. Leveraging existing technologies such as GitHub and CircleCI and extensively using *containers* (Docker and Singularity), the proposed ecosystem automatically generates the appropriate container images with a minimized impact on the researcher’s development flow. This paper shows how these apps are created, stored, and executed either locally or in HPCs. To maximize interoperability and reduce manual metadata handling, BIDS Apps require that input data are in the BIDS organization format. The ultimate goal of BIDS Apps is reproducibility, thus this framework is particularly focused on archiving and versioning of neuroimaging workflows.

## Materials and Methods

### Engaging the community

To kickstart the repository of BIDS Apps, a four-day long coding sprint was organized at Stanford University in August 2016. Leading neuroimaging methods and workflow developers were invited to learn about BIDS, Docker, and Singularity. In addition, the sprint was advertised on the Stanford Center for Reproducible Neuroscience and Twitter, and outreach was performed during the 2016 OHBM Meeting in Geneva. The workshop consisted of one day of training; during the remaining three days, hands-on support was provided by experienced Docker developers.

### Command-line specification

**Figure 2.**
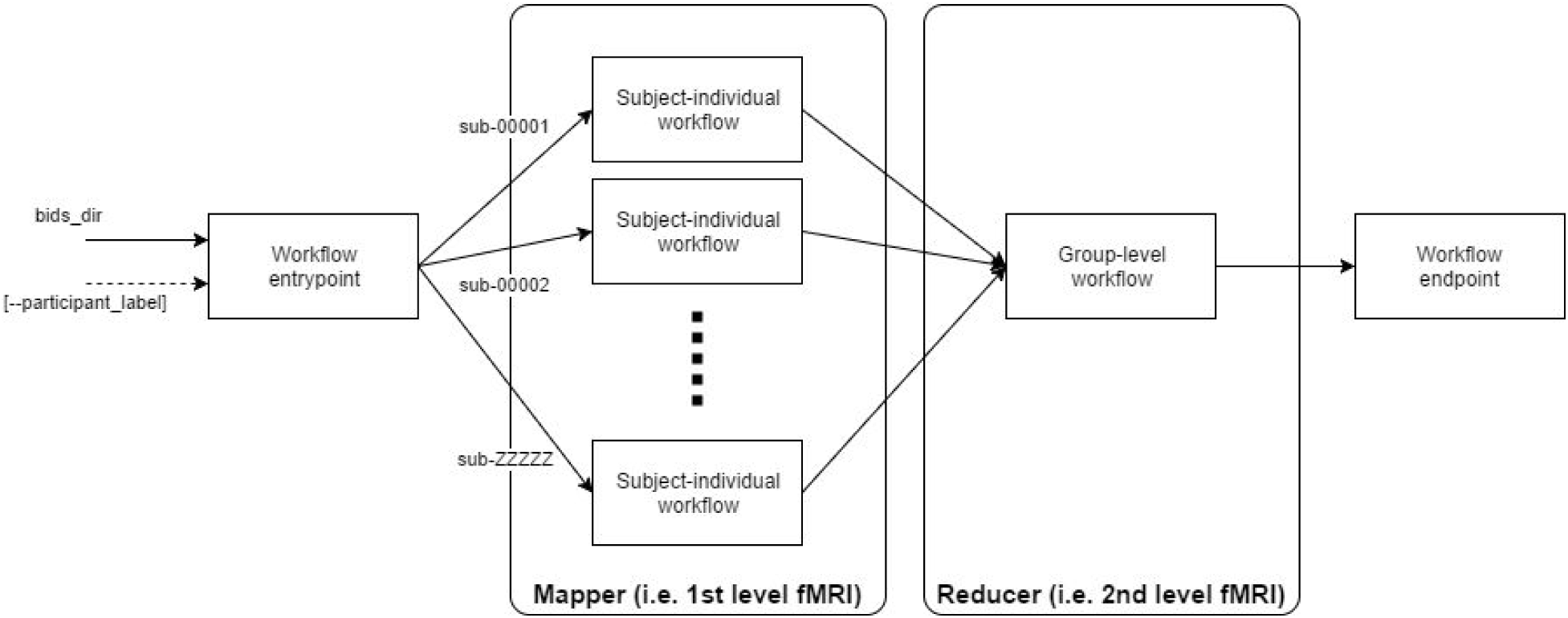
**The overall structure of workflows:** workflows will need to decouple the individual level analysis (process independent subjects) from the group-level analysis. For the analysis of individual subjects, the workflow will require an understanding of the <u>BIDS structure</u> so that the required inputs for the designated subject are found. The optional reducer module will take up from results generated in the mapper and generate a group output. The overall workflow has an entrypoint and an endpoint responsible of setting-up the map-reduce tasks and the tear-down including organizing the outputs for its archiving, respectively. Please mind that each app can implement multiple map and reduce steps (see Advanced use cases).

This approach to run our workflows requires sticking with three *standards*: 1) a common command-line interface, 2) a Docker container to ensure portability, and 3) a standard for organizing input data. Containers created this way can be easily integrated into OpenfMRI as well as other data analysis platforms. Thanks to Docker-to-Singularity conversion they can also be easily run on High Performance Computers (clusters) without the need to install all of the dependencies.

Each workflow/pipeline will be run independently for each subject (the map step). Results of this execution (arranged in whatever way the pipeline prefers) can be optionally processed in a group level analysis (reduce step).

### Command line interface

Each pipeline should have a simple wrapper script used to run it. The script should be a command-line interface and accept the following command-line arguments (minimally):

1. For the participant level (aka map) step:
  - `bids_dir` - (positional argument #1) the directory with the input dataset formatted according to the BIDS standard. This directory is read only.
  - `output_dir` - (positional_argument #2) the directory where the output files should be stored. This is the only directory the pipeline should write to. Can be used to store intermediate files, but they should be removed after the pipeline finishes. This directory is shared across all of the participant level jobs - it’s up to the script to create subfolders for each subject.
  - `“participant”` - (positional_argument #3) indicates that this is a participant level analysis.
  - `--Participant_label` - (optional) label of the participant that should be analyzed. The label corresponds to sub-<participant_label> from the BIDS spec (so it does not include “sub-”). If this parameter is not provided all subjects should be analyzed. Multiple participants can be specified with a space separated list. Example:
    Run processing for every subject independently (map step). Each of these operations can be performed in parallel. There are no restrictions or specification of how data should be organized inside the `output_dir`.
    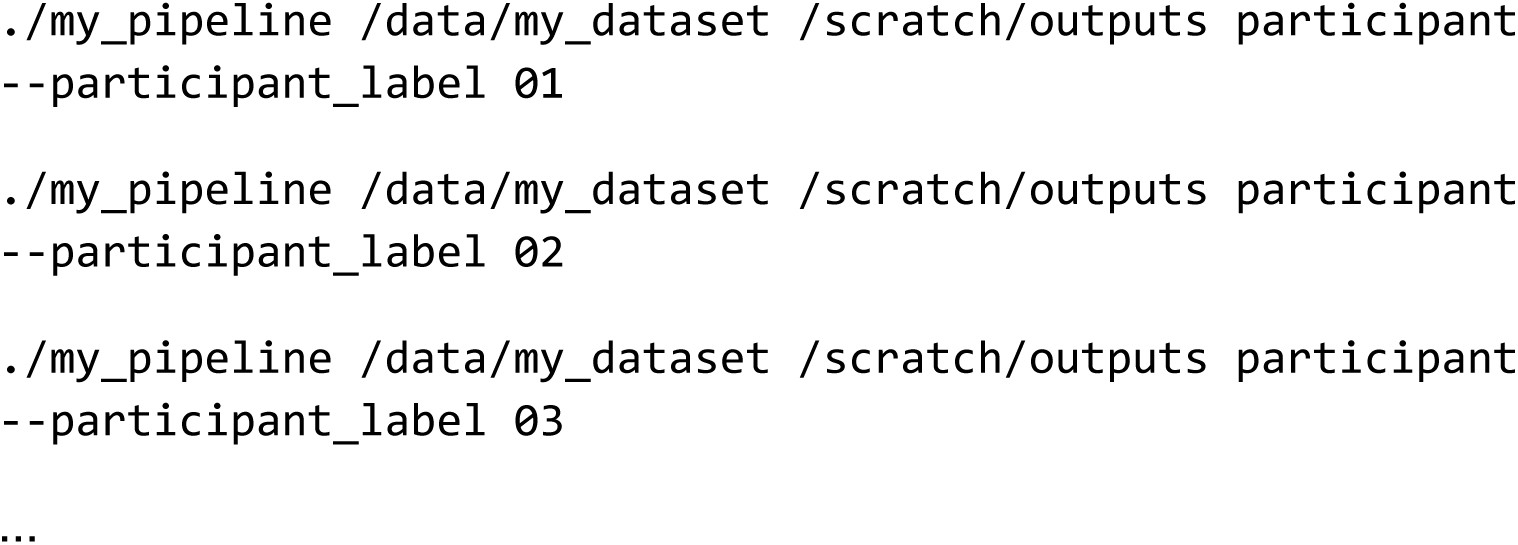
2. (Optional) For the group level (aka reduce) step:
  - `bids_dir` - (positional argument #1) the directory with the input dataset formatted according to the BIDS standard. This directory is read only.
  - `output_dir` - (positional_argument #2) the directory where the output files should be stored. This is the only directory the pipeline should write to. Can be used to store intermediate files, but they should be removed after the pipeline finishes. This directory is the same one that was used in the participant level analysis and should be prepopulated with participant level results before running the group level.
  - “group” - (positional_argument #3) indicates that this is a group level analysis.
  - `--Participant_label` - (optional) labels of the participants that should be analyzed. The label corresponds to sub-<participant_label> from the BIDS spec (so it does not include “sub-”). If this parameter is not provided all subjects should be analyzed. Multiple participants can be specified with a space separated list. This can be useful if you want to do a group level analysis on a subset of all participants in your dataset Example: ~~~
./my_pipeline /data/my_dataset /scratch/outputs group
~~~

The script can also accept other arguments specific to your pipeline (see for example`--template_name` in <u>FreeSurfer App</u>). Mind that the same set of extra arguments will be passed to the map (single subject level) and the reduce (group level) stage.

### Advanced use cases

- Multiple map reduce steps. In case your pipeline needs to do multiple map reduce steps the analysis_level (third positional argument) can take additional arguments: `participant2, group2, participant3, group3` etc. In the description of your app please specify how many map reduce steps are necessary.
- Within-job multi-CPU parallelization. Each job (on any given level: participant or group) might be run on a multi-CPU machine; in such case, a `parameter--n_cpus` followed by an integer will be passed with the number of CPUs available.
- If your app is capable of adapting its workflow depending on how much memory is available on the environment it is running on you can implement an optional `--mem_mb` flag. When running your app the execution system will pass available memory in megabytes.

## References

1 Hanke M, Halchenko YO. Neuroscience runs on GNU / Linux. Front Neuroinform. 2011;5: 7–9. doi: 10.3389/fninf.2011.00008

2 Halchenko YO, Hanke M. Open is not enough. Let’ s take the next step: An integrated, community-driven computing platform for neuroscience. Front Neuroinform. 2012;6. doi: 10.3389/fninf.2012.00022

3 Boekel W, Wagenmakers E-J, Belay L, Verhagen J, Brown S, Forstmann BU. A purely confirmatory replication study of structural brain-behavior correlations. Cortex. 2015;66: 115–133. doi: 10.1016/j.cortex.2014.11.019

4 Mackenzie-Graham AJ, Van Horn JD, Woods RP, Crawford KL, Toga AW. Provenance in neuroimaging. Neuroimage. 2008;42: 178–195. doi: 10.1016/j.neuroimage.2008.04.186

5 Gronenschild EHBM, Habets P, Jacobs HIL, Mengelers R, Rozendaal N, van Os J, et al The Effects of FreeSurfer Version, Workstation Type, and Macintosh Operating System Version on Anatomical Volume and Cortical Thickness Measurements. Hayasaka S, editor. PLoS One. 2012;7: e38234. doi: 10.1371/journal.pone.0038234

6 Glatard T, Lewis LB, da Silva RF, Adalat R, Beck N, Lepage C, et al Reproducibility of neuroimaging analyses across operating systems. Front Neuroinform. Frontiers; 2015;9. doi: 10.3389/fninf.2015.00012

7 Angiuoli SV, Matalka M, Gussman A, Galens K, Vangala M, Riley DR, et al CloVR: a virtual machine for automated and portable sequence analysis from the desktop using cloud computing. BMC Bioinformatics. 2011;12: 356. doi: 10.1186/1471-2105-12-356

8 Krampis K, Booth T, Chapman B, Tiwari B, Bicak M, Field D, et al Cloud BioLinux: pre-configured and on-demand bioinformatics computing for the genomics community. BMC Bioinformatics. 2012;13: 42. doi: 10.1186/1471-2105-13-42

9 Folarin AA, Dobson RJB, Newhouse SJ. NGSeasy: a next generation sequencing pipeline in Docker containers. F1000Res. 2015;4. doi: 10.12688/f1000research.7104.1

10 Moreews F, Sallou O, Ménager H, Le Bras Y, Monjeaud C, Blanchet C, et al BioShaDock: a community driven bioinformatics shared Docker-based tools registry. F1000Res. 2015;4: 1443. doi: 10.12688/f1000research.7536.1

11 Devisetty UK, Kennedy K, Sarando P, Merchant N, Lyons E., Bringing your tools to CyVerse Discovery Environment using Docker. F1000Res. 2016;5. doi: 10.12688/f1000research.8935.1

12 Belmann P, Dröge J, Bremges A, McHardy AC, Sczyrba A, Barton MD. Bioboxes: standardised containers for interchangeable bioinformatics software. Gigascience. 2015;4: 47. doi: 10.1186/s13742-015-0087-0

13 Jordan C, Skidmore E, Dooley R. The iPlant collaborative: cyberinfrastructure for plant biology. Front Plant Sci. journal.frontiersin.org; 2011; Available: http://journal.frontiersin.org/article/10.3389/fpls.2011.00034/full.

14 Kurtzer GM. Singularity 2.1.2 - Linux application and environment containers for science [Internet]. 2016. doi: 10.5281/zenodo.60736

15 Gorgolewski KJ, Auer T, Calhoun VD, Craddock RC, Das S, Duff EP, et al The brain imaging data structure, a format for organizing and describing outputs of neuroimaging experiments. Sci Data. 2016; 3: 160044. doi: 10.1038/sdata.2016.44

16 Gorgolewski K, Burns CD, Madison C, Clark D, Halchenko YO, Waskom ML, et al Nipype: a flexible, lightweight and extensible neuroimaging data processing framework in python. Front Neuroinform. 2011;5: 13. doi: 10.3389/fninf.2011.00013

17 Dean J, Ghemawat S. MapReduce: simplified data processing on large clusters. Commun ACM. 2008;51: 107. doi: 10.1145/1327452.1327492

18 Gorgolewski KJ, Storkey A, Bastin ME, Whittle IR, Wardlaw JM, Pernet CR. A test-retest fMRI dataset for motor, language and spatial attention functions. Gigascience. 2013;2: 6. doi: 10.1186/2047-217X-2-6

19 Fischl B. FreeSurfer Neuroimage 2012; 62 774–781. doi: 10.1016/j.neuroimage.2012.01.021

20 Reuter M, Schmansky NJ, Rosas HD, Fischl B., Within-Subject Template Estimation for Unbiased Longitudinal Image Analysis. Neuroimage. 2012;61: 1402–1418. doi: 10.1016/j.neuroimage.2012.02.084

21 Kiar G, Gray Roncal W, Mhembere D, Bridgeford E, Burns R, Vogelstein JT. ndmg: NeuroData’s MRI Graphs pipeline [Internet]. Zenodo; 2016. doi: 10.5281/zenodo.60206

22 Kiar G, Gorgolewski KJ, Kleissas D, Roncal WG, Litt B, Wandell B, et al Science In the Cloud (SIC): A use case in MRI Connectomics [Internet]. arXiv [q-bio.QM]. 2016. Available: http://arxiv.org/abs/1610.08484.

23 Eklund A, Dufort P, Villani M, Laconte S. BROCCOLI: Software for fast fMRI analysis on many-core CPUs and GPUs. Front Neuroinform. 2014;8: 24. doi: 10.3389/fninf.2014.00024

24 Raffelt DA, Smith RE, Ridgway GR, Tournier J-D, Vaughan DN, Rose S, et al Connectivity-based fixel enhancement: Whole-brain statistical analysis of diffusion MRI measures in the presence of crossing fibres. Neuroimage. 2015;117: 40–55. doi: 10.1016/j.neuroimage.2015.05.039

25 Raffelt DA, Tournier J-D, Smith RE, Vaughan DN, Jackson G, Ridgway GR, et al Investigating White Matter Fibre Density and Morphology using Fixel-Based Analysis. Neuroimage. 2016; doi: 10.1016/j.neuroimage.2016.09.029

26 Friston KJ. Statistical Parametric Mapping: The Analysis of Functional Brain Images [Internet]. Elsevier/Academic Press; 2007. p. 647. Available: http://books.google.com/books?id=KZZcLqkxhFIC.

27 Zarrar S, Steven G, Qingyang L, Yassine B, Chaogan Y, Zhen Y, et al The Preprocessed Connectomes Project Quality Assessment Protocol - a resource for measuring the quality of MRI data. Front Neurosci. 2015;9. doi: 10.3389/conf.fnins.2015.91.00047

28 Craddock C, Sikka S, Cheung B, Khanuja R, Ghosh SS, Yan C, et al Towards Automated Analysis of Connectomes: The Configurable Pipeline for the Analysis of Connectomes (C-PAC). Front Neuroinform. doi: 10.3389/conf.fninf.2013.09.00042

29 Guntupalli JS, Hanke M, Halchenko YO, Connolly AC, Ramadge PJ, Haxby JV. A, Model of Representational Spaces in Human Cortex. Cereb Cortex. 2016;26: 2919–2934. doi: 10.1093/cercor/bhw068

30 Klein A, Tourville J. 101 labeled brain images and a consistent human cortical labeling protocol. Front Neurosci. 2012;6: 171. doi: 10.3389/fnins.2012.00171

31 Smith RE, Tournier J-D, Calamante F, Connelly A., The effects of SIFT on the reproducibility and biological accuracy of the structural connectome. Neuroimage. 2015;104: 253–265. doi: 10.1016/j.neuroimage.2014.10.004

32 Abraham A, Pedregosa F, Eickenberg M, Gervais P, Mueller A, Kossaifi J, et al Machine learning for neuroimaging with scikit-learn. Front Neuroinform. 2014;8: 14. doi: 10.3389/fninf.2014.00014

33 Cusack R, Vicente-Grabovetsky A, Mitchell DJ, Wild CJ, Auer T, Linke AC, et al Automatic analysis (aa): efficient neuroimaging workflows and parallel processing using Matlab and XML. Front Neuroinform. 2014;8: 90. doi: 10.3389/fninf.2014.00090

34 Bellec P, Lavoie-Courchesne S, Dickinson P, Lerch JP, Zijdenbos AP, Evans AC., The pipeline system for Octave and Matlab (PSOM): a lightweight scripting framework and execution engine for scientific workflows. Front Neuroinform. 2012;6: 7. doi: 10.3389/fninf.2012.00007

35 Glasser MF, Sotiropoulos SN, Wilson JA, Coalson TS, Fischl B, Andersson JL, et al The minimal preprocessing pipelines for the Human Connectome Project. Neuroimage. 2013;80: 105–124. doi: 10.1016/j.neuroimage.2013.04.127

36 Smith SM, Andersson J, Auerbach EJ, Beckmann CF, Bijsterbosch J, Douaud G, et al Resting-state fMRI in the Human Connectome Project. Neuroimage. 2013; doi: 10.1016/j.neuroimage.2013.05.039

37 Chen P-H (cameron), Chen J, Yeshurun Y, Hasson U, Haxby J, Ramadge PJ. A Reduced-Dimension fMRI Shared Response Model. Advances in Neural Information Processing Systems 28. Curran Associates, Inc. 2015. pp. 460–468. Available: http://papers.nips.cc/paper/5855-a-reduced-dimension-fmri-shared-response-model.pdf.

38 Strother SC, Anderson J, Hansen LK, Kjems U, Kustra R, Sidtis J, et al The quantitative evaluation of functional neuroimaging experiments: the NPAIRS data analysis framework. Neuroimage. 2002;15: 747–771. doi: 10.1006/nimg.2001.1034

39 Churchill NW, Oder A, Abdi H, Tam F, Lee W, Thomas C, et al Optimizing preprocessing and analysis pipelines for single-subject fMRI. I. Standard temporal motion and physiological noise correction methods. Hum Brain Mapp. 2012;33: 609–627. doi: 10.1002/hbm.21238

40 Churchill NW, Yourganov G, Oder A, Tam F, Graham SJ, Strother SC. Optimizing preprocessing and analysis pipelines for single-subject fMRI: 2. Interactions with ICA, PCA, task contrast and inter-subject heterogeneity. PLoS One. 2012;7: e31147. doi: 10.1371/journal.pone.0031147

41 Churchill NW, Spring R, Afshin-Pour B, Dong F, Strother SC., An Automated Adaptive Framework for Optimizing Preprocessing Pipelines in Task-Based Functional MRI PLoS One 2015 10 e0131520 doi: 10.1371/journal.pone.0131520

42 Chakravarty MM, Rapoport JL, Giedd JN, Raznahan A, Shaw P, Collins DL, et al Striatal shape abnormalities as novel neurodevelopmental endophenotypes in schizophrenia: a longitudinal study. Hum Brain Mapp. 2015;36: 1458–1469. doi: 10.1002/hbm.22715

43 Pipitone J, Park MTM, Winterburn J, Lett TA, Lerch JP, Pruessner JC, et al Multi-atlas segmentation of the whole hippocampus and subfields using multiple automatically generated templates. Neuroimage. 2014;101: 494–512. doi: 10.1016/j.neuroimage.2014.04.054

44 Park MTM, Pipitone J, Baer LH, Winterburn JL, Shah Y, Chavez S, et al Derivation of high-resolution MRI atlases of the human cerebellum at 3T and segmentation using multiple automatically generated templates. Neuroimage. 2014;95: 217–231. doi: 10.1016/j.neuroimage.2014.03.037

45 Chakravarty MM, Steadman P, Eede MC, Calcott RD, Gu V, van Shaw P, et al Performing label-fusion-based segmentation using multiple automatically generated templates. Hum Brain Mapp. 2013;34: 2635–2654. doi: 10.1002/hbm.22092

46 Yendiki A, Panneck P, Srinivasan P, Stevens A, Zöllei L, Augustinack J, et al Automated probabilistic reconstruction of white-matter pathways in health and disease using an atlas of the underlying anatomy. Front Neuroinform. 2011;5: 23. doi: 10.3389/fninf.2011.00023

47 Di Tommaso P, Palumbo E, Chatzou M, Prieto P, Heuer ML, Notredame C., The impact of Docker containers on the performance of genomic pipelines. PeerJ. 2015;3: e1273. doi: 10.7717/peerj.1273

48 Marcus DS, Olsen TR, Ramaratnam M, Buckner RL. The extensible neuroimaging archive toolkit. Neuroinformatics. Humana Press; 2007;5: 11–33. doi: 10.1385/NI:5:1:11

49 Sherif T, Rioux P, Rousseau M-E, Kassis N, Beck N, Adalat R, et al CBRAIN: a web-based, distributed computing platform for collaborative neuroimaging research. Front Neuroinform. 2014;8: 54. doi: 10.3389/fninf.2014.00054

50 Glatard T, da Silva RF, Boujelben N, Adalat R, Beck N, da Rioux P, et al Boutiques: an application-sharing system based on Linux containers. Neuroinformatics 2015. doi: 10.3389/conf.fnins.2015.91.00012

51 Hale JS, Li L, Richardson CN, Wells GN. Containers for portable, productive and performant scientific computing [Internet]. arXiv [cs.DC]. 2016. Available: http://arxiv.org/abs/1608.07573.

47 Hale JS, Li L, Richardson CN, Wells GN. Containers for portable, productive and performant scientific computing [Internet]. arXiv [cs.DC]. 2016. Available: http://arxiv.org/abs/1608.07573.

